# Social relationships, social isolation, and the human gut microbiota

**DOI:** 10.1101/428938

**Authors:** Kimberly A. Dill-McFarland, Zheng-Zheng Tang, Julia H. Kemis, Robert L. Kerby, Guanhua Chen, Alberto Palloni, Thomas Sorenson, Federico E. Rey, Pamela Herd

**Affiliations:** Department of Bacteriology, U. of Wisconsin-Madison, 1550 Linden Drive, Madison, WI, USA 53706; Center for the Demography of Health and Aging, 1180 Observatory Drive, Madison, WI, USA, 53706; Department of Biostatistics and Medical Informatics, U. of Wisconsin-Madison, 600 Highland Avenue, Madison, WI, USA, 53792; Wisconsin Institute for Discovery, 330 N Orchard St, Madison, WI, USA, 53715; Department of Sociology, U. of Wisconsin-Madison, 1180 Observatory Drive, Madison, WI, USA, 53706

**Author notes:** Current-address: Department of Microbiology and Immunology, U. of British Columbia, 2350 Health Sciences Mall, Vancouver, BC, Canada, V6T 1Z3. Equal contribution (Kimberly A. Dill-McFarland, Zheng-Zheng Tang). F. E. Rey 5157 Microbial Sciences, 1550 Linden Drive, Madison, WI, USA 53706. P. Herd 3454 Social Science, 1180 Observatory Drive, Madison, WI, USA, 53706.

## Abstract

Social relationships shape human health and mortality via behavioral, psychosocial, and physiological mechanisms, including inflammatory and immune responses. Though not tested in human studies, recent primate studies indicate that the gut microbiome may also be a biological mechanism linking relationships to health. Integrating microbiota data into the 60-year-old Wisconsin Longitudinal Study, we found that socialness with family and friends is associated with differences in the human fecal microbiota. Analysis of spouse (N = 94) and sibling pairs (N = 83) further revealed that spouses have more similar microbiota and more bacterial taxa in common than siblings, with no observed differences between sibling and unrelated pairs. These differences held even after accounting for dietary factors. The differences between unrelated individuals and married couples was driven entirely by couples who reported close relationships; there were no differences in similarity between couples reporting somewhat close relationships and unrelated individuals. Moreover, the microbiota of married individuals, compared to those living alone, has greater diversity and richness, with the greatest diversity among couples reporting close relationships, which is notable given decades of research documenting the health benefits of marriage. These results suggest that human interactions, especially sustained, close marital relationships, influence the gut microbiota.

## INTRODUCTION

Social relationships exert a sustained influence on human health and mortality with social isolation having strong negative consequences and high levels of social integration far exceeding the protective effects on mortality of individual level behaviors such as smoking cessation or maintaining a normal weight ^1,2^. Research in the social sciences has shown that individuals who cohabitate in marriage and marital like relationships have better health than do unpartnered adults^3^. For both social relationships generally, and marriage specifically, health benefits are largely achieved in the context of high-quality relationships. The robust links between these relationships and health are related to stress, behaviors, and psychosocial resources, among other factors ^2^. In part, social support may impact one’s health by reinforcing healthy habits, reducing the impacts of stress, and preventing the use of unhealthy “self-medications” like smoking and drinking ^2^. Additional research points to stress-related biological processes that may also contribute to the positive impacts of social relationships through changes in inflammatory processes, metabolic syndrome, and neurological functioning ^4,5^.

Recent work in the field of microbiology points to another possible biological mechanism linking human relationships and health: the microbiome. The microbial communities that inhabit mammals have profound effects on biology and health ^6^. Gastrointestinal (GI) microbial communities impact host health by modulating the epigenome ^7^, brain function ^8^, and metabolism of drugs and nutrients ^9^ as well as impacting immune system function ^10^ and development ^11^. While the microbiota reaches an adult-like configuration by three to five years of age ^12^, considerable variation exists between adults ^13^, and differences are mediated by a number of factors. Most notable among these are diet ^14^ and host genetics ^15^, which also correlate with health. An individual’s microbiota structure (*i.e.* relative abundance) and composition (*i.e.* who’s there) can change rapidly in response to inputs like diet ^16^ and antibiotics ^17^. Nonetheless, there is evidence that an individual’s microbiota remains relatively stable over many years ^18–20^, perhaps in part because a person’s behaviors also tend to be consistent over many years.

While a number of factors like diet are known to impact both the microbiota and health ^21^, less is known regarding social relationships. Most existing research has focused on animal models, which has produced compelling evidence that social interactions, via a range of different types of physical contact, influences the gut microbiota through microbial sharing between individuals ^22–26^. Additionally, states of isolation, such as maternal neglect, influence the gut microbial composition in animal models ^27^ at least in part through stress ^28,29^. Thus, the gut microbiota may play a role in some of the long-term health effects of social relationships.

But despite this tantalizing evidence, studies in human populations remain relatively small in number ^30^. There are a few studies exploring how mother-infant interactions influence the development of the infant’s gut microbiome and even how broader social interactions influence the milk microbiome ^31,32^. In terms of adults, there is evidence regarding the influence of cohabitation, may influence the gut microbiome. A few recent studies have found that individuals living together had more similar gut ^33^ and skin ^33,34^ microbiota. Interestingly, however, another study found that married cohabitating couples had no more similarity in the composition of their gut microbiota than did unrelated individuals ^35^.

Thus, while it does appear that living together may influence the gut microbiome, human studies have not investigated how adult relationships, rather than just simply living in the same space, may influence the gut microbiome. The quality of the relationship may matter. Closer relationships likely lead to even closer shared environments, via mechanisms such as time spent physically together. Indeed, one recent study of wild baboons found that close partners within social groups had more similar gut microbiotas ^36^. Studies have also have not more generally compared how living alone versus living with an intimate partner influences the gut microbiome; individuals living alone are on average, de facto, more socially isolated than those living with someone, and animal studies have generally shown that social isolation leads to decreased microbial diversity ^22,37–39^. Though causality is not certain, decreased microbial diversity is associated with obesity, cardiac disease, and type 2 diabetes, and a range of other inflammatory disorders ^40–47^. More broadly, there is extensive evidence that cohabitating couples in later life have substantially improved physical and psychological well-being compared to single adults ^48–50^. Thus, similar mechanisms might explain some of the variance in findings in humans.

An important hindrance to research examining social relationships and the GI microbiota is the availability of human samples with sufficiently well-characterized life course measures of broader social environments and conditions. Thus, most microbiological research in this field is based on animal models ^22–25^. However, there are now a wide array of well-characterized longitudinal studies in the social sciences that have generated decades of research documenting relationships between broader social environments and mortality ^5,51–54^. These data can provide a platform for studies of the human microbiota to advance knowledge for both social scientists and microbiologists, including whether social conditions influence the gut microbiota and whether the gut microbiota is a mediating biological mechanism explaining how social conditions influence health.

Here, we leverage a multidisciplinary collaboration to investigate the links between human interaction, the microbiota, and human health. We utilized data in the nearly 60-year Wisconsin Longitudinal Study (WLS) ^54^, which constitutes a random sample of 1 in 3 1957 Wisconsin high school graduates (N = 10,317), as well as selected spouses and siblings surveyed periodically during their adult life. We correlate the fecal microbiota of 408 older individuals (58 – 91 yo) from WLS with extensive health and behavioral data, as well as compare spouse and sibling pairs within the cohort. Overall, this project demonstrates the promise of joint participation between social scientists and microbiologists in efforts to more fully understand the gut microbiota and its impacts on human health.

## RESULTS and DISCUSSION

We employed 16S rRNA gene sequencing to characterize the fecal microbiota of 408 individuals, including Wisconsin Longitudinal Study (WLS) graduates (N = 179, 76 ± 0.5 years old), siblings of graduates (134, 74 ± 6.4), spouses of graduates (63, 76 ± 3.7), and spouses of siblings (32, 73 ± 6.1). We then correlated these communities to longitudinal survey data collected from 1957 to 2015 as part of WLS ^54^. For more details on this data collection, see ^55^. A total of 24.5 million high-quality sequences were obtained for 408 fecal samples (60,000 ± 19,000 SD sequences per sample) after quality filtering in mothur. All samples achieved sufficient coverage as determined by Good’s coverage > 99% (Dataset S1).

In the WLS graduate cohort, we identified several factors correlated with gastrointestinal (GI) microbiota including sex, antibiotics, dietary protein, high blood sugar, and heart disease (Fig. 1, Fig. S1, Table S1). These factors were reported in the previous literature ^57,58^ with diet playing a particularly strong role ^14,16,56^. Thus, we assessed diet across a number of measures including habitual intake of protein, vegetables, and fruits (Text S1) during the year prior to the fecal sample collection (for details, see METHODS, Statistical analysis for graduates). While overall dietary dissimilarity (Bray-Curtis and Jaccard) across these three categories correlated with gut microbiota dissimilarity, only the total frequency of dietary protein consumption was robustly associated with microbial composition using either univariate or multivariate analyses (Table S1). Thus, we note that all analyses have adjusted for potential confounders including age, sex, antibiotics, dietary protein, and chronic conditions (diabetes and heart disease) unless stated otherwise. In some analyses ‐that are noted below—we do also include vegetable and fruit dietary data.

**Figure 1.**
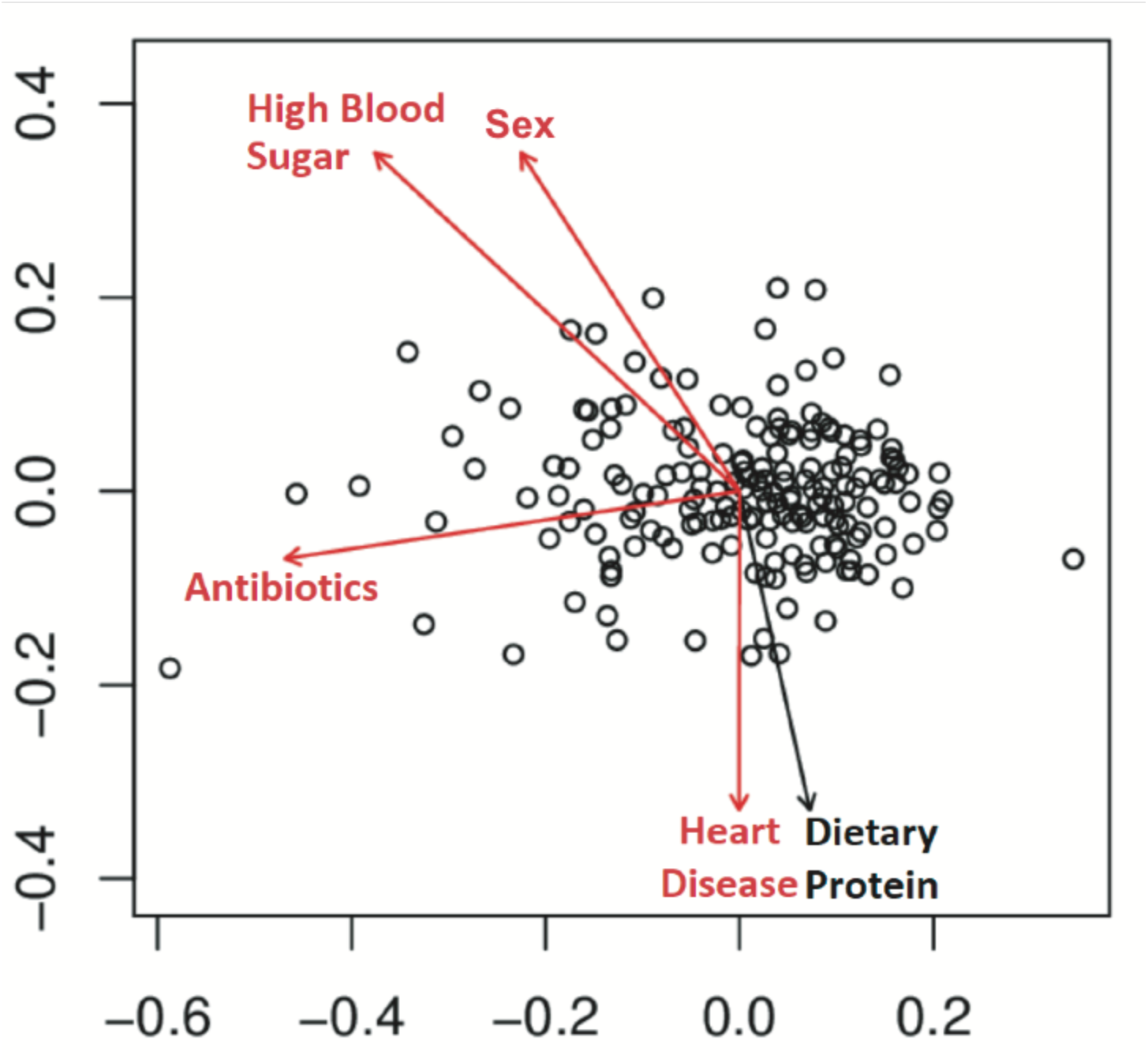
Factors associated with the overall fecal microbiota. Non-metric multidimensional scaling (nMDS) of unweighted UniFrac for all graduates (N = 179). Variables found to be significant (PERMANOVA P < 0.05, red) and trends (0.05 < P < 0.1, black) are shown as fitted arrows. Arrows point toward increasing values (dietary protein), toward affirmative responses (high blood sugar, antibiotics, heart disease), or from male to female (sex).

### Social interactions and the human fecal microbiota

Human interactions were also associated with differences in gut microbiota and diversity. Specifically, we found that individuals that were cohabitating with a spouse or partner had more similar microbiota composition with their cohabitating spouse/partner as well as higher diversity and richness than unmarried, non-cohabitating individuals (unweighted UniFrac *P* = 0.029 Shannon *P* = 0.005, Chao *P* = 0.011, Fig. 2). Since all cohabitating pairs were male-female and sex was a strong determinant of the microbiota in this study (*P* < 0.001, Table S2), increased diversity may be partially due to sustained exchange of microorganisms between the sexes, though we were not able to test this given that there were no same sex couples in these data. Increases in diversity seen here are consistent with a previous cohabitation study in pigs ^59^ and may have implications for human health, as previous work indicates that increased gut microbial diversity is associated with lower risks of irritable bowel syndrome (IBS), Crohn’s disease, ulcerative colitis, and other GI afflictions ^60^.

**Figure 2.**
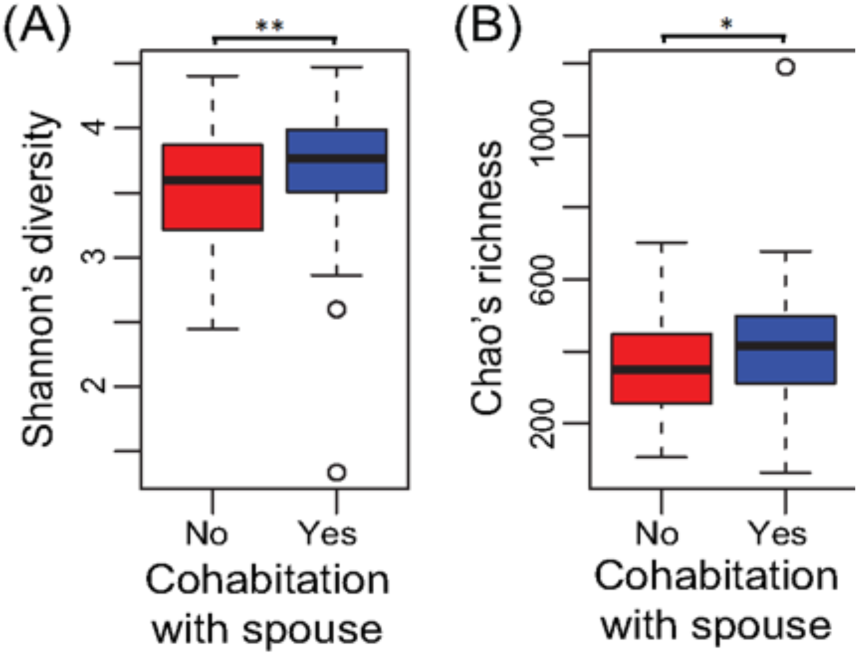
Cohabitation is associated with increased alpha-diversity. Boxplots of (A) Shannon’s diversity and (B) Chao’s richness of graduates that are (blue) or are not (red) cohabitating with a spouse or partner. All spouses/partners were cohabitating while all non-cohabitating individuals were unmarried. **P < 0.01, *P < 0.05

Social interactions with relatives and friends were stronger predictors of gut microbial diversity in non-cohabiting individuals than cohabiting spouses/partners (unweighted UniFrac *P* = 0.0030, Shannon *P* = 0.042, Chao *P* = 0.063, Fig. S2) (Table S2). Here, social interactions were defined as the sum of “How many times during the past four weeks have you gotten together with relatives/friends?” The associations may have been weaker for cohabitating spouses due to their higher microbial diversity; ecological theory supports that diverse communities are more resilient and resistant to invasion by new species ^61^. Thus, one explanation for these differential associations is that the more diverse microbiotas of individuals already cohabitating with a spouse may not have been as strongly influenced by increasing social interactions while the less diverse microbiotas of those living alone were more strongly influenced by invasion of new species through social exposures. It is also possible that cohabitating couples share the same friends and socialize together with these friends. However, factors contributing to the resilience of the human gut microbiota require further exploration to confirm this hypothesis.

### Spouses have more similar microbes than siblings and unrelated individuals

Previous studies have established that the GI microbiota reaches an adult-like configuration by 3 to 5 years of age ^18,62,63^ and that during adulthood, communities are stable on the time scale of years ^19,20^. Thus, microbial communities established in early life may persist and, aside from extreme perturbation, remain stable across one’s adult lifetime. However, our analyses comparing sibling, couple, and unrelated pairs challenge the assumption that microbial communities established in early life will be largely unperturbed in later life (for details, see METHODS, Statistical Analysis for spouse and siblings). In fact, we find no evidence for a remaining influence of early life on the composition of the gut microbiota among older adults. In this older cohort, spouses were more similar than unrelated subjects (unweighted UniFrac *P* = 3.2E-5) or sibling pairs (unweighted UniFrac *P* = 0.033, Fig. 3). Further, the length of the cohabitating marital relationship was positively correlated with similarity (unweighted UniFrac, *P* = 0.031)In contrast, siblings were no more similar than unrelated pairs by any beta-diversity metric (*P* > 0.3, Fig. 3A, Fig. S3A, D, G) (Table S2). We also found no evidence that the physical proximity of siblings—as measured by physical distance between siblings—influenced gut microbial similarity. Thus, adult factors like marriage with cohabitation (spouses) appear to have a greater influence on the adult gut microbiota than early-life environment or genetics (siblings).

**Figure 3.**
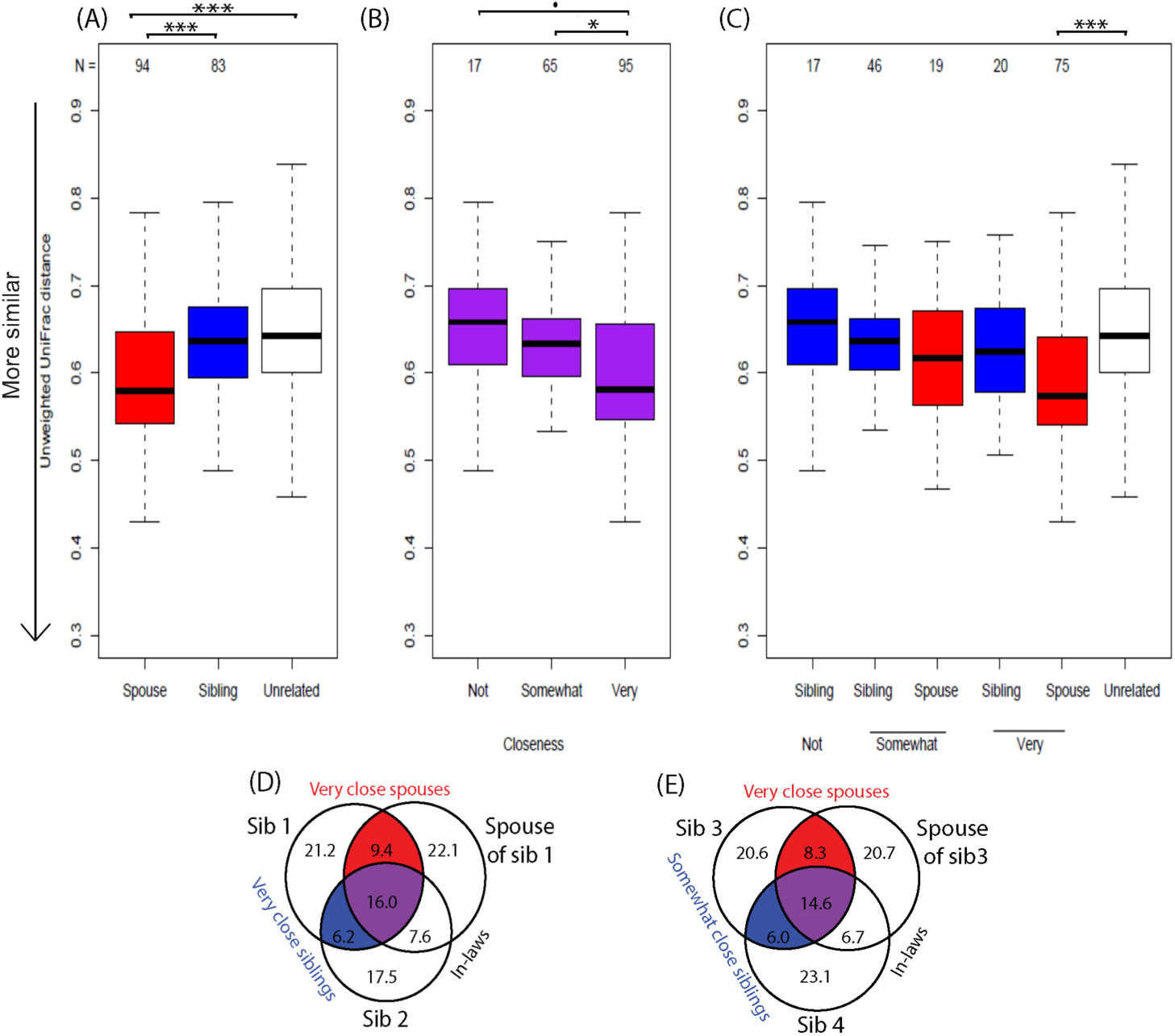
Microbial sharing in spouse and sibling relationships. Unweighted UniFrac distances of (A) spouse, sibling, and unrelated pairs, (B) spouses and siblings grouped by relationship closeness, and (C) spouses and siblings separated by relationship closeness. Groups (A) were compared in linear regression model adjusting for potential confounders (*e.g.* age, sex, diet, health conditions). P-values were averaged across 1000 rounds of unrelated pair sampling. Closeness groups (B,C) were compared in linear regression models adjusting for potential confounders. (D, E) Average percentages of shared OTUs within family groups including a related spouse and sibling pair. Families included those with both very close spouses and siblings (D, N = 12) and those with very close spouses and somewhat close siblings (E, N = 17). Percentages are of the total number of OTUs across all three individuals, and circle sizes are proportional to total percentages represented. ***P < 0.001, **P < 0.01, *P < 0.05, •P< 0.1

This is further supported by our findings that childhood farm status was not associated with microbial richness (Chao *P* = 0.342) while working on a farm as an adult correlated with higher richness (Chao *P* = 0.005). Farm-driven differences in the microbiota are of particular interest, because adolescents that grew up on a farm have more diverse microbial communities ^64^ and reduced risk of asthma and other atopic diseases both during childhood ^65^ and as adults ^66^. Given the results here, it appears that the microbially-driven protective effects of early farm exposures are not due to the persistence of protective microorganisms acquired in early-life. Protection may, instead, be conferred by immune development and training by early-life microbes as suggested previously ^67^.

Our results are also in contrast with previous work showing that genetically related individuals harbor more similar microbial communities than unrelated individuals, regardless of current cohabitation ^35,68–70^. However, these previous studies investigated children ^68^, young adults ^35,69^, or a wide age range ^70^, and therefore, cumulative changes across a lifetime may not have reached a level sufficient to overcome early-life factors impacting the microbiota. Additionally, sibling pairs in other studies were twins ^35,68–70^, and many focused on monozygotic twins (same sex and age) ^35,69,70^ as opposed to this study where siblings were often of opposite sexes (43%) and ranged from less than a year to 18 years apart in age. Also, the unrelated group in this study may have exhibited higher homogeneity than unrelated groups in other studies, because most grew up in and/or currently live in the state of Wisconsin. Thus, compared to previous studies, siblings were likely less similar and unrelated pairs more similar across our cohort. Furthermore, genetic effects on the microbiota are often small ^70^ and detection may require a larger human cohort than used here. Taken together, these factors may have contributed to the lack of significant differences observed between sibling and unrelated groups even though average sibling beta-diversity was intermediate between spouses and unrelated individuals.

### Increased microbial similarity, diversity, and richness in closer relationships

For both spouse and sibling relationships, microbiota similarity was associated with self-reported relationship closeness (unweighted UniFrac *P* = 0.0079). Closeness was measured by participant responses to “How close are you and your current spouse/sibling?” on a scale of not at all (1) to very (4). Due to the small sample sizes in the categories “Not very” (N=13) and “Not at all” (N=4), we combined these two groups into “Not” close. Across spouses and siblings, individuals in very close relationships harbored gut microbial communities more similar to their close social partners than those in not very close relationships (Fig. 3B), though this relationship was not significant within the spousal and sibling pair groups separately (Fig. 3C). Moreover, differences between spouses and unrelated individuals, in terms of closeness (Fig. 2), as well as the enhanced diversity and richness in cohabitating couples versus individuals living alone (Fig. 2) were driven by spouses reporting very close relationships. This was in contrast to couples reporting only somewhat close relationships as these pairs did not have higher gut microbiota similarity than unrelated pairs (Table S3) nor did they display microbial diversity or richness different from non-cohabitating individuals (Table S3). Importantly, the apparent impacts of relationship closeness do not appear to be mediated by similarities in diet since overall dietary dissimilarity (Bray-Curtis and Jaccard) did not significantly differ according to relationship closeness (ANOVA *P* > 0.5; Table S4). We note that these included sensitivity tests that modeled diet based on the protein consumption, but also overall diet that captured vegetable and fruit consumption.

While diet is often correlated with the GI microbiota ^56^, closeness points to the less well-understood contributions of human interactions and shared behaviors. Close proximity and frequent physical contact were correlated with microbiota similarity among primates with direct microbial sharing between individuals contributing to similarity ^22,23^. In this study, relationship closeness may represent a summative measure of time spend together, physical affection, and other human interactions with the potential to result in microbial sharing. Indeed, there is evidence that the salivary microbiome influences the gut microbiome and the salivary microbiome may be influenced by kissing ^71,72^. In these data, this is supported by the fact that spouses had more operational taxonomic units (OTUs, a proxy for microbial species) in common (30.4 ± 7.32%) than siblings (26.4 ± 7.47%, t-test *P* = 4.39E-04) (Dataset S2). Also, when comparing the spouse and sibling pair within a family represented in this dataset, a person tended to have more OTUs in common with his or her very close spouse (25.4 ± 7.9%) than his or her very close sibling (22.2 ± 6.4%, N = 12 families, *P* = 0.074, Fig. 3D). This is also true when comparing very close spouses (22.9 ± 5.8%) and somewhat close siblings within a family (20.6 ± 5.5%, N = 17 families, *P* = 0.027, Fig. 3E).

### Shared taxa with close human relationships

In general, highly abundant genera and OTUs were shared between many spouse and sibling pairs while less abundant shared taxa were specific to one pair type and shared by a small number of pairs within that type (Dataset S3). OTUs that were commonly found among spouses or siblings (> 50% of pairs) but rare in the unrelated dataset (< 70% individuals, < 49% unrelated pairs) may represent bacterial species easily shared by close human interaction. These OTUs were predominately from the phylum Firmicutes (16 of 22 OTUs) with representatives of families Lachnospiraceae and Ruminococcaceae (Dataset S4). Interestingly, most of these potentially shared OTUs were from strictly anaerobic taxa, indicating that persisting in an oxygen-rich environment in-between hosts may not be a limiting factor in very close human relationships. Transmission, in these cases, could be mediated by direct contact similar to mechanisms of vertical transmission from mother to child ^73^.

Taxa commonly associated with reduced disease incidence or severity like *Akkermansia muciniphila* ^74^, *Bifidobacterium* spp. ^75,76^, *Collinsella aerofaciens* ^76^, and *Ruminococcus bromii* ^77^ as well as potentially harmful taxa like *Clostridium spiroforme* ^78,79^ were often present in both persons in a spouse or sibling pair. Several of these potentially shared OTUs were associated with disease incidence in the larger dataset. In particular, *Ruminococcus bromii*, *Lachnospira* spp. and unclassified Ruminococcaceae and Lachnospiraceae OTUs were less abundant in those with high blood sugar (Fig. 4, Dataset S4). These results are in contrast to previous reports of more abundant Ruminococcaceae/*Ruminococcus* ^80,81^ and Lachnospiraceae ^80^ associated with diabetes in humans and may point to important differences in the impacts of the microbiota on metabolic health in older populations. Overall, though, this indicates that GI microbial species with the potential to impact host health may be shared by close human interactions. However, it cannot be discounted that these apparent health associations may be mediated by diet as those with high blood sugar often consume specific diets to manage disease.

**Figure 4.**
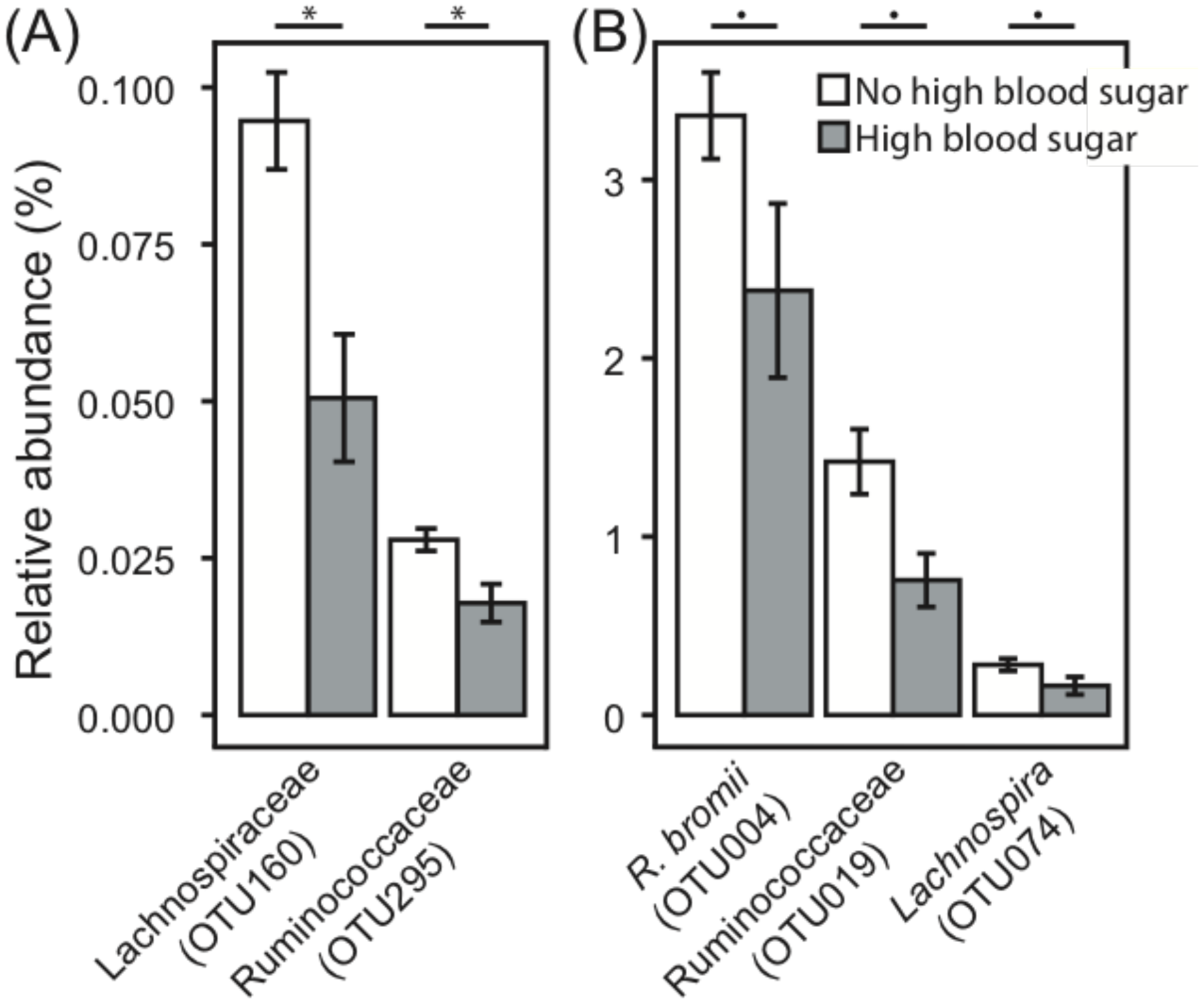
Percent relative abundance of OTUs that are commonly shared between spouses and that differed between those with (grey) and without (white) high blood sugar. (A) Low abundance and (B) more highly abundant OTUs. Means with standard error bars are shown. Kruskal-Wallis FDR *P < 0.05, •P < 0.1

Overall, our findings indicate that in order to understand environmental influences on the gut microbioata, we must now consider the many microbiotas with which this individual interacts. Socialness with family and friends is associated with differences in the fecal microbiota. These differences held even after accounting for dietary factors, though given this is the first study of its kind, it will be critical for future work to validate this finding. Thus, it is possible that relationships with others may influence the gut microbiota and consequent health outcomes, either through direct microbial transfer or reinforcement of healthy microbiota behaviors. We further found not only that married couples had more similar gut microbiota but also that the microbiota of married individuals, compared to those living alone, has greater diversity and richness. Key to both of these findings, however, was that they were driven by individuals reporting that they were very close to their spouse as opposed to somewhat close. Close marriage relationships had a stronger influence than the shared genetic factors and early life environments among siblings. This finding is interesting, in part, because it parallels an extensive body of evidence demonstrating robust links between high quality marriages and morbidity and mortality. Future work could attempt to disentangle the mechanisms linking close relationships to microbial composition. For example, while we did not find evidence that shared diet was primarily responsible for these findings, we could not test precise frequencies of physical contact and intimacy as an alternative explanatory mechanism. Importantly, the types of physical contact and intimacy change over the life course, with sexual intimacy becoming far less frequent in later life, but other kinds of intimate physical contact remaining important. Regardless of the mediating mechanism, from a social and population health science perspective, decades of evidence that social relationships, especially close ones like marriage, influence morbidity and mortality make the central finding of significant interest. For example, even if future work finds a greater role for shared diets, it is still the social relationships that drive that shared diet. Overall, these results provide support for the gut microbiome as a possible mediating pathway between social relationships, especially marriage, and health and mortality. These findings, in the context of the robust body of evidence linking social relationships to human morbidity and mortality, provide fodder for further work examining the role of the gut microbiome as a possible biological mediator in these relationships ^32^. Further microbiota work across time in a more diverse population should be undertaken with the many longitudinal social science studies currently underway in an effort to increase our understanding of the complex interactions between human behavior, the microbiota, and health.

## METHODS

### Wisconsin Longitudinal Study (WLS)

WLS is based on a one-third sample of all 1957 Wisconsin high school graduates (N = 10,317) as well as selected siblings and spouses ^54^. Graduates originally enrolled with an in-person questionnaire upon graduating high school in 1957, which was followed by data collection in 1964, 1975, 1992, 2004, and 2011. Siblings were surveyed in 1977, 1994, 2005, and 2011; spouses were surveyed in 2004 or 2006. The content of WLS surveys changed to reflect the participants’ life course with an education focus in the initial data collection, familial and career outcomes in young adulthood / midlife, and health, cognitive functioning, psychological well-being, non-work activities, caregiving, bereavement, social support, and end-of-life preparations in later rounds. WLS data collection was approved by the Institutional Review Board (IRB) at the University of Wisconsin-Madison (2014-1066, 2015-0955). Informed consent, the content and procedures of which were included in the IRB approval, was obtained from participants. All methods were performed in accordance with relevant guidelines and regulations.

### Study design

A total of 500 individuals were randomly drawn from the full WLS dataset constrained based on the following: 1) participated in the 2011 interviews; 2) lived in one of 10 counties in Wisconsin that included both northern rural counties and southern more urban counties; and 3) were part of a sibling pair. Individuals were removed from the study if they did not give consent, their sample did not arrive for processing chilled, but not frozen, within 48 hrs of collection, or their sample did not yield at least 10,000 sequences for analysis. This resulted in 408 individuals being included in this study.

An additional survey was administered at the time of fecal sampling, which detailed dietary data from the prior three days, prescription/antibiotic use, current living situation, and additional health information. This as well as selected data from the larger WLS study focused on health, spouse/sibling relationships, and social interactions were used in this study (Text S1). Data, documentation, and other materials are accessible at http://www.ssc.wisc.edu/wlsresearch. Access to the full dataset can be obtained through.

### Sample collection

Stool samples were collected by participants in November 2014, January 2015, or April 2015 following provided instructions (Text S2). Participants stored samples at ∼4 °C in their refrigerator or in a NanoCool box (Albuquerque, NM) with cooling cartridge and customized foam insert, supplemented with a single ice pack. Interviewers picked-up samples from participants within 24 hours of collection and shipped samples in fresh NanoCool boxes for arrival at UW-Madison within 48 hours of collection. Upon arrival, an aliquot of feces was collected for DNA extraction and immediately stored at −80°C until further processing. The use of WLS and fecal microbiota data were approved by the Institutional Review Board at the University of Wisconsin-Madison (2017-0600).

### DNA extraction

Genomic DNA was extracted from fecal aliquots using a bead-beating protocol

^45^. Briefly, feces (∼100 mg) were re-suspended in a solution containing 500 μl of extraction buffer [200 mM Tris (pH 8.0), 200 mM NaCL, 20 mM EDTA], 210 μl of 20% SDS, 500 μl phenol:chloroform:isoamyl alcohol (pH 7.9, 25:24:1) and 500 μl of 0.1-mm diameter zirconia/silica beads. Samples were mechanically disrupted using a bead beater (BioSpec Products, Barlesville, OK; maximum setting for 3 min at room temperature), followed by centrifugation, recovery of the aqueous phase, and precipitation with isopropanol. QIAquick 96-well PCR Purification Kit (Qiagen, Germantown, MD) was used to remove contaminants. Isolated DNA was eluted in 5 mM Tris/HCl (pH 8.5) and was stored at −80 °C until further use. We also note that we used negative controls.

### Sequencing

PCR was performed using universal primers flanking the variable 4 (V4) region of the bacterial 16S rRNA gene ^82^. We used negative controls for each PCR reaction. PCR reactions where the negative control yielded a product were not sequenced until the problem was solved. Samples were processed all together, not in batches, in a random order (i.e., not clustered by family). Additionally, unlike other specimens (e.g., saliva, skin), DNA contamination from reagents is in general not a problem for fecal samples given the high DNA content of the sample (10^12^ microbes/g of feces). In one reaction per sample, 10 - 50 ng DNA, 10 μM each primer, 12.5 μl 2X HotStart ReadyMix (KAPA Biosystems, Wilmington, MA, USA), and water to 25 μl were used. Cycling conditions were initial denaturation of 95 °C for 3 min followed by 25 cycles of 95 °C for 30 s, 55 °C for 30 s, and 72 °C for 30 s, with a final extension of 72 °C for 5 min. PCR products were purified with the QIAquick 96-well PCR Purification Kit (Qiagen, Germantown, MD, USA). Samples were quantified by Qubit Fluorometer (Invitrogen, Carlsbad, CA, USA) and equimolar pooled. The pool plus 5% PhiX control DNA was sequenced through the U. of Wisconsin-Madison Biotechnology Center with the MiSeq 2×250 v2 kit (Illumina, San Diego, CA, USA) using custom sequencing primers ^82^. All DNA sequences are available upon institutional review board (IRB) or other ethics board approval through.

### Sequence clean-up

All sequences were demultiplexed on the Illumina MiSeq. Sequence cleanup and processing was performed with mothur v.1.36.1 ^83^ following a protocol similar to ^82^. Briefly, paired-end sequences were combined into contigs with default parameters (match bonus = 1, mismatch penalty = −1, gap penalty = −2, gap extend penalty = −1, insert quality 20, mismatch quality difference 6). Poor-quality sequences, including those with ambiguous base pairs, homopolymers greater than 8, or outside 200 – 500 bp in length, were discarded. Sequences were then aligned to the SILVA 16S rRNA gene reference alignment database ^84^ and trimmed to the V4 region. To reduce sequencing error, sequences with 2 or fewer differences were pre-clustered. Chimera detection and removal were performed using UCHIME ^85^. Final sequences were then classified to the GreenGenes database ^86^. Singletons were removed to facilitate downstream analyses. All sequences were grouped into 98% operational taxonomic units (OTUs) by uncorrected pairwise distances and average neighbor clustering in mothur. Clustering performed on uncorrected pairwise distances revealed no differences in clusters at 97 vs 98% similarity. Therefore, the stricter cutoff was reported Coverage was assessed by Good’s coverage, and then samples were normalized to whole number counts by percent relative abundance to approximately 10,000 sequences per sample (9,914 - 10,061 after rounding.

### Statistical analysis for graduates

Graduates were assessed separately from siblings and spouses to avoid potential interactions, and the graduate subset was not significantly different from other groups (PERMANOVA *P* Bray-Curtis *P* = 0.56, Jaccard *P* = 0.57, weighted UniFrac *P* = 0.33, unweighted UniFrac *P* = 0.24). Alpha-diversity was assessed with Shannon’s diversity and Chao’s richness calculated in mothur. Differences in alpha-metrics were assessed in R v3.3.2 ^87^ by linear regression with the Benjamini-Hochberg correction for multiple comparisons across each metric.

Microbial beta-diversity was assessed for Bray-Curtis, Jaccard, weighted, and unweighted UniFrac metrics with results shown for unweighted UniFrac unless otherwise noted. Dietary beta-diversity was assessed for Bray-Curtis and Jaccard metrics as well as corresponding nMDS axes calculated from habitual intake of specific sources of protein (N = 4), vegetables (N = 76), and fruits (N = 24) expressed as times consumed per week (protein), proportions of total types (all), and presence/absence of individual types (all). Differences in beta-diversity were tested with permutational analysis of variance (PERMANOVA, adonis) in the vegan package ^88^ with the Benjamini-Hochberg correction for multiple comparisons across each metric and a maximum of 5000 permutations. All variables were modeled using independent, univariate tests and dietary variables were additionally modeled using multivariate tests of all components (protein, vegetables, fruits). Co-variance of microbial and dietary beta metrics was measured using Mantel’s test. The factors that associated with the microbiome in univariate models (*i.e.* age, sex, antibiotics, dietary protein, high blood sugar, and heart disease ^57^) were adjusted for in regression models as potential confounders. Beta-diversity was visualized by non-metric multidimensional scaling (nMDS) plots with arrows from significant variables (PERMANOVA) fitted to the ordination using maximum correlation (envfit, vegan). All tests were assessed at significance *P* < 0.05 and trends 0.05 < *P* < 0.1.

### Statistical analysis for spouses and siblings

For the spouse and sibling similarity analysis, the unit of the observation is the pair (*i.e.* spouse, sibling, or unrelated pair defined below) and the variables used in the analysis are distance in individual measurements between the two members of the pair. Specifically, beta-diversity metrics were used to quantify the distance in microbial and overall diet whereas absolute difference were calculated to quantify the distance in all the other variables (*e.g.* age, sex, dietary protein). We sampled unrelated pairs from the data in order to compare the spouse or sibling pair with unrelated pairs. In particular, the unrelated individuals cannot be siblings, spouses, or in-laws, and each unrelated pair will match the corresponding spouse or sibling pair in sex and antibiotics usage. Beta-diversity distances were compared among spouse, sibling, and unrelated pairs using linear regression while adjusting for the distance in age, sex, dietary protein, health conditions (if available). P-values were averaged across 1000 rounds of unrelated pair sampling. For closeness analysis, we removed age and sex from the model because the two variables are highly correlated with pair type (*i.e.* sibling/spouse pair can be accurately classified using the difference of the age or sex between the two members of the pair). For comparing OTU sharing among spouse and sibling within a family, we used mixed-effect models to account for family clustering. All tests were assessed at significance *P* < 0.05 and trends 0.05 < *P* < 0.1.

## AUTHOR CONTRIBUTIONS

FER and PH designed research. JHK, RLK, and TS completed sample processing and sequencing. AP contributed to the conceptual and analytic approach to conducting the analyses. KADM, ZZT, and GC analyzed the data. KADM, ZZT, GC, FER, and PH contributed to the final manuscript. Alberto Palloni^5^

## ACKNOWLEDGMENTS

The authors wish to thank members of the Rey laboratory for their insights and support. This work was supported by the National Institute of Food and Agriculture, U.S. Department of Agriculture (2016-67017-24416, FER), the National Institutes for Aging, National Institutes for Health (AG041868, PH), the Center for the Demography of Health and Aging (AG017266, PH), and the Clinical and Translational Science Award (CTSA), National Institutes for Health National Center for Advancing Translational Sciences (NCATS) (UL1TR000427, ZZT). Additional support was provided by the Vice Chancellor for Research and Graduate Education (VCGRE) at the U. of Wisconsin-Madison. The content is solely the responsibility of the authors and does not necessarily represent the official views of the NIH.

The authors declare that we have no competing interests as defined by Nature Research, or other interests that might be perceived to influence the results and/or discussion reported in this paper.

